# Ubiquinone is a hysteretic modulator of the NADH:cytochrome *b*_5_ reductase activity of human C*b*_5_R

**DOI:** 10.64898/2026.01.14.699433

**Authors:** Gabriel N. Valerio, Oscar H. Martinez-Costa, Carlos Sanchez-Cabeza, Cristina M. Cordas, Alejandro K. Samhan-Arias

## Abstract

**Background:** Cytochrome *b*_5_ reductase is a flavoprotein that transfers electrons from NADH to multiple electron acceptors, such as cytochrome *b*_5_ or ubiquinone. Hysteresis is a phenomenon characterized by a slow transition between active and inactive catalytic states, leading to a lag phase in enzymatic activity. In this study, the effect of the soluble analogue of ubiquinone named 2,3 dimethoxy-5-methyl-1,4 benzoquinone (CoQ_0_) on the NADH:C*b*_5_ reductase activity of recombinant human soluble C*b*_5_R, using recombinant human soluble C*b*_5_ as a substrate was evaluated. The aim of this study was to determine whether ubiquinone exerts a hysteretic modulation of this activity based on previous studies supporting that microsomal reduction of cytochrome *b*_5_ is controlled by redox hysteresis.

**Results:** The NADH:cytochrome *b*_5_ reductase activity of C*b*_5_R was characterized at different concentrations of C*b*_5_R, cytochrome *b*_5_, and CoQ_0_ by monitoring the reduction of cytochrome *b*_5_. The addition of CoQ_0_ induced the appearance of a lag phase, whose duration increased with the concentration of CoQ_0_ and decreased with higher concentrations of cytochrome *b*_5_ or C*b*_5_R. Additionally, a concentration-dependent decrease in the maximum rate of reduction and the appearance of positive cooperativity was observed in the presence of CoQ_0_ which resulted in leading to lower *K*_M_ values for cytochrome *b*_5_. This suggests the formation of a CoQ_0_:C*b*_5_R complex altering the interaction between the reductase and cytochrome *b*^5^ which increases the affinity for cytochrome *b*_5_. Cyclic voltammetry data support the formation of CoQ_0_/protein complex that could be responsible for the hysteretic behavior.

**Conclusions:** These results support the hypothesis that CoQ_0_ is a hysteretic modulator and inhibitor of the NADH: cytochrome *b*_5_ reductase activity of human C*b*_5_R.

## INTRODUCTION

A hysteretic behavior has been described for many enzymes in the literature, including the pyruvate phosphate dikinase (PPDK) from *Trypanosoma cruzi* [1], glutamate dehydrogenase from *Neurospora crassa* [2], and wheat germ hexokinase [3]. The importance of hysteresis in the regulation of certain NADH-dependent enzymes and complexes, such as complex I of the electron transport chain, that exhibits hysteretic modulation by ubiquinone (CoQ_10_), has also been reported [4]. Given that complex I is capable of catalyzing the reduction of CoQ_10_ coupled to NADH oxidation [5–7], it is reasonable to question whether some other non-mitochondrial enzymes, that consume NADH to reduce CoQ_10_ and other substrates, could be hysteretically modulated by quinones.

Cytochrome *b*_5_ (C*b*_5_) is a haemprotein electron carrier and cofactor in multiple enzymatic reactions, such as those catalyzed by monooxygenases, dioxygenases, desaturases [8], by donating electrons to multiple electron acceptors ( e.g. methaemoglobin [9], desaturases [10] and P450s [11]). Two main enzymes can reduce C*b*_5_ in biological membranes: the NADPH-cytochrome P450 reductase (P450R) and the NADH-cytochrome *b*_5_ reductase (C*b*_5_R). The levels of C*b*_5_R are key for some of the reactions occurring in microsomes and a regulation of some P450 has been found to be mediated by the C*b*_5_/C*b*_5_R system [12].

Regarding C*b*_5_R, two main forms of the enzyme exist: a membrane-bound and a soluble form, sharing a common water-soluble domain and differ by hydrophobic segments present in the membrane form that anchor the protein to membranes [13]. The NADH-C*b*_5_R contains a flavin adenine dinucleotide (FAD) binding domain and a nicotinamide adenine dinucleotide (phosphate) binding domain. In terms of electron transfer mechanism, we have recently reviewed the mechanism of electron transfer of NADH-C*b*_5_R to electron acceptors [13]. The FAD cofactor is critical for the enzyme’s electron transfer activity, as it accepts electrons from NADH and transfers them in one- or two-electron reactions [13]. The electron transfer reactions require the formation of a complex with NADH that is oxidized during the catalytic reactions. The oxidation of NADH by the flavoprotein occurs in less than 2ms and at least one intermediate is involved in the flavin reduction and one in the flavin reoxidation, complete the oxidative cycle. Oxidation of the reduced flavoprotein occurs faster than the reduction of the enzyme which is important for the observed turnover. In one electron reduction/oxidation reactions, a flavin semiquinone radical can be generated which has been shown to be formed independently of the substrate used (C*b*_5_ or ferricyanide). In the final step, NAD^+^ slowly auto-oxidizes and disproportionate the FAD semiquinone [14]. One electron can react with NAD^+^, yielding NAD^•^ [15]. Also, one electron can react with the NAD^+^-bound oxidized enzyme yielding the blue and red semiquinone (or a mixture of the two forms of the enzyme).

Ubiquinone (CoQ_10_) is a lipid-soluble compound crucial for cellular energy production. It features a quinone head group and an isoprenoid side chain, rendering it hydrophobic and enabling its integration into lipid membranes. CoQ_10_ is an essential component of the mitochondrial redox chain, shuttling electrons from Complex I to Complex III which is key for establishing the proton gradient necessary for ATP synthesis during cellular respiration [16]. An alteration of mitochondrial respiration leads to free radical production due to pro-oxidant properties of CoQ_10_ [17,18]. Nevertheless, a protective function against oxidative stress has been suggested for the extramitochondrial pool of this molecule [19]. Reduced levels of CoQ_10_ have been postulated as key for the protection against lipid peroxidation and the control of cellular processes like ferroptosis [20]. Therefore, those systems in charge of reducing CoQ_10_ are emerging as potential candidates for the control of the cellular fate [20]. We believe that the new dual functions as antioxidant and pro-oxidant of CoQ_10_, as redox mediators controlling redox sensitive pathways, are important to be explored.

C*b*_5_R is one of the extramitochondrial NADH-dependent enzymes with the ability to reduce CoQ_10_. It presents the potential to be hysteretically modulated by CoQ_10,_ as observed in mitochondrial complex I [4,21]. We have described that the microsomal NADH-dependent reduction activity of C*b*_5,_ to which C*b*_5_R is one of the main contributors, shows hysteretic modulation by CoQ_10_ [22]. The main objective of the current study is to experimentally characterize the effect of CoQ_0_ using the soluble form of CoQ_10_ on this enzymatic activity measured with recombinant human soluble C*b*_5_R and soluble C*b*_5_. Kinetic profiles displaying lag phases are expected, as well as modifications in kinetic parameters dependent on the concentration of the added modulator, and consistent with hysteresis.

## MATERIALS AND METHODS

### Expression and purification of recombinant human soluble C*b*_5_

Chemical-competent BL21 *E. coli* strains were transformed with the Heb5 plasmid, containing the codifying sequence for soluble C*b*_5_, and plated on LB agar with 0.1 mg/ml ampicillin as described in [23,24]. One colony was picked up and grown for 8 h at 37 °C in 5 mL of LB media supplemented with ampicillin. Then, the culture was transferred into 1 L of Terrific Broth media supplemented by ampicillin and grown for 24 h at 37 °C. After 24 hours, the speed was reduced, and the culture was maintained under shaking for 12 hours more. Cells were pelleted after overgrowth culture centrifugation at 6000 rpm for 30 min.

For protein purification, pellets were incubated in 50 mM Tris, 1 mM EDTA, 1 mM PMSF pH 8.1, and 0.5 mg/mL lysozyme for 1 h. After lysis, 50 mM of MgCl_2_, 0.2% deoxyribonuclease, 0.5% Triton X-100, 0.5M NaCl, and 0.05 mg/ml of DNase were added, and the lysate was left under agitation for 1 h. The soluble fraction was separated from debris by centrifugation at 9,000*g* using a JA20 rotor for 30 min at 4 °C. The sample was precipitated with ammonium sulphate up to 50% saturation, was put to agitate for 1 hour, and then centrifuged at 8000g.The supernatant was extensively dialyzed against 10 mM Tris, 1 mM EDTA pH 8.1 at 4 °C and loaded onto a diethylamino ethyl (DEAE) Sepharose column (2.5 × 30 cm) (GE Healthcare) previously equilibrated in 10 mM Tris, 1 mM EDTA pH 8.1. A gradient was performed stepwise by the addition of increasing concentrations of Tris buffer (up to 200 mM) to the buffer in the presence of 1 mM EDTA, 0.5% Tx-100, and 0.2% deoxycholate. C*b*_5_ was eluted in 10 mM Tris, 1 mM EDTA, and 250 mM potassium thiocyanate pH 8.1. Eluent was cleaned from contaminants by the addition of ammonium sulphate 1.1 M and passing it through a CL Sepharose 4B column (2.5×10 cm) (equilibrated with 100 mM Tris, 1 mM EDTA) with no retention. After concentration with an Amicon filter (10-kDa cutoff), the almost pure protein was further separated from contaminants using a Sephadex G75 column (GE Healthcare) (2.5×50 cm) equilibrated with Tris 150 mM pH 7.5. After elution, C*b*_5_ was concentrated with an Amicon filter (10 kDa cutoff) and 30% glycerol was added before freezing at −80 °C.

For protein quantification, the absorbance change at 557 nm was registered using a Shimadzu UV-mini 1240 spectrophotometer and a 10mm quartz cuvette filled with 200 times diluted C*b*_5_ in phosphate buffer 20 mM at pH 7.0. C*b*_5_ was reduced by addition of a few crystals of dithionite. A differential extinction coefficient of 16.5 mM^-1^.cm^-1^ was used for reduced-oxidized C*b*_5_ (15).

### Expression and purification of recombinant human soluble C*b*_5_R

Expression and purification of recombinant human soluble C*b*_5_R was performed as shown in [23]. Quantification of the activity was performed by measuring the NADH: ferricyanide reductase activity at 25°C using potassium ferricyanide (0.5 mM) and NADH (1 mM) presence of C*b*_5_R in potassium phosphate 20 mM, DTPA 0.1 mM, pH 7.4 using a Shimadzu UVmini-1240 spectrophotometer and a quartz cuvette with 10mm bandpass. For quantification an extinction coefficient for ferricyanide at 420 nm of 1mM^-1^.cm^-1^ was used [25]. For protein normalization a enzymatic value for this activity of 270µmol/min/mg of C*b*_5_R was used [23].

### Measurement of the NADH:CoQ_0_ reductase activity of C*b*_5_R

The NADH:CoQ_0_ reductase activity of C*b*_5_R was measured in potassium phosphate 20 mM, DTPA 0.1 mM, pH 7.0 after addition of NADH 500 µM at the concentration CoQ_0_ and C*b*_5_R indicated in the text, at 25 °C using a Perkin Elmer Lambda 40 spectrophotometer and a quartz cuvette with 10 mm bandpass, following the decrease in absorbance at 410 nm and an extinction coefficient for CoQ_0_ at this wavelength of 0.7 mM^-1^.cm^-1^ [26].

### Measurement of the NADH:C*b*_5_ reductase activity of C*b*_5_R

The NADH:C*b*_5_ reductase activity of C*b*_5_R in was measured in potassium phosphate 20 mM, DTPA 0.1 mM, pH 7.0 after addition of NADH 150 µM at the concentration C*b*_5_ and C*b*_5_R indicated in the text, at 25 °C using a (Perkin Elmer Lambda 40 spectrophotometer and a quartz cuvette with 10 mm bandpass, following the decrease in absorbance at 557 nm and an extinction coefficient for CoQ_0_ at this wavelength of 16.5 mM^-1^.cm^-1^ [25]. The study of CoQ_0_ effect on this activity was assayed by adding the CoQ_0_ concentrations indicated in the text (4 mM stock solution prepared on phosphate buffer 20 mM pH 7.0).

### Measurement of the NADH oxidation

The NADH consumption in the assays was measured in the presence of C*b*_5_R (50 ng/mL) in potassium phosphate 20 mM, DTPA 0.1 mM, pH 7.0 in the absence of presence of C*b*_5_ (30 µM) and/or CoQ_0_ (5 µM), after addition of NADH 50 µM to the assay. The absorbance was measured at 340 nm using Shimadzu UVmini-1240 spectrophotometer and a quartz cuvette with 10mm bandpass. For NADH quantification an extinction coefficient of 6.22 mM^-1^.cm^-1^ at 340 nm was used. For quantification of NADH consumption in assays performed in the presence of C*b*_5_, the recorded absorbance at 340 nm was corrected by that of C*b*_5_ reduction at this wavelength by subtraction of this contribution using a differential extinction coefficient for reduced minus oxidized C*b*_5_ at 340 nm of 9.2 mM^-1^.cm^-1^ calculated for this study.

### Docking

#### Protein Preparation

The three-dimensional structure of cytochrome *b*₅ reductase (C*b*_5_R) was obtained from the Protein Data Bank (PDB ID: 1UMK, which corresponds to the human erythrocyte C*b*_5_R). Protein preparation was carried out using UCSF Chimera’s Dock Prep [27]. All crystallographic water molecules, ions, and non-essential ligands were removed, while the FAD cofactor was retained due to its functional relevance. Hydrogen atoms were added according to default Chimera parameters, and the structure was optimized to relieve steric clashes. Partial atomic charges for the FAD cofactor were assigned automatically by Chimera during Dock Prep according to Gasteiger [28], resulting in a net charge of −1 for the cofactor.

#### Ligand Preparation

The CoQ₀ ligand was constructed using Avogadro 1.2.0 software [29] and energy-minimized by applying the steepest descent algorithm, General AMBER force field (GAFF) [28]. Dock prep was performed in UCSF Chimera 1.16. Partial atomic charges were assigned according to Gasteiger [28].

#### Docking Procedure

Molecular docking simulations were performed using AutoDock Vina 1.1.2 [30,31]. The docking grid was set to involve the whole protein (x, y and z coordinates for the grid center: 54.9776, 39.6201, −1.0365; x, y and z coordinates for the size: 47.6992, 46.1357, 46.933). Independent docking runs were performed with an exhaustiveness value of 8, each producing ten binding poses, yielding a total of 100 docking poses. The results were compiled with their respective AutoDock Vina binding affinity scores and RMSD’s. Molecular graphics and analyses performed with UCSF Chimera, developed by the Resource for Biocomputing, Visualization, and Informatics at the University of California, San Francisco, with support from NIH P41-GM103311. For defining the binding site, common residues present in the top ranked poses in the triplicate experiments were used as performed in other publications, and the binding energy was defined as the average of the calculated for top ranked poses in triplicate experiments [32,33].

### Electrochemistry

Electrochemical assays were performed inside a Faraday Cage. An Autolab PGSTAT 12 Potentiostat/Galvanostat was used to acquire the signals. A 3 electrodes one compartment cell setup was used, where platinum was the counter electrode, pyrolytic graphite was the working electrode, and a saturated calomel electrode (SCE) was used as reference. The electrochemical cell setup was deaerated with a continuous flow of argon for 15 min. The working electrode was pre-treated by polishing with 2 grades of alumina (1 and 0.3 μm, in this order) subjected to a water ultrasound bath for 3 min and washed thoroughly with Mili-Q water. The working electrode was then dried with an argon flow and 5 µL of samples were placed on the working electrode’s surfaces. After a decrease in the initial volume to half (partial solvent casting), a 3.5 kDa cut-off membrane was used to entrap the proteins. The CoQ_0_ was soluble in the cell electrolyte. The formal potential calculation was calculated considering the average value of the corresponding redox process. Other parameters, such as the number of electrons associated to a process, are calculated using different equations accordingly to if the species are in solution or physical entrapped in the membrane, where in this case, a thin-layer model was used [34].

### Data analysis

The data shown in this work are the result of average of replicate experiments (minimum in triplicate), and the estimated error corresponds to the calculation of the standard deviation associated with those values. The fitting of the data to nonlinear regression models was carried out using the Solver tool in Microsoft Excel.

## RESULTS

### Determination of kinetic parameters of the NADH:CoQ_0_ reductase activity of C*b*_5_R

NADH:CoQ_0_ reductase activity of C*b*_5_R was first characterized to obtain the kinetic parameters of this enzymatic reaction. First, we analyzed the dependence of the activity with the concentration of C*b*_5_R. The reduction of CoQ_0_ (1 mM) was monitored by analyzing the decrease in absorbance at 410 nm, in the presence of NADH (0.5 mM) with increasing concentration of C*b*_5_R, namely, 1, 2.5 and 5 µg/mL (Figure 1a). The dependence of the CoQ_0_ reduction rate on the enzyme concentration fitted with a linear regression with a R^2^ = 0.99 with slope of 17.1 ± 0.4 μmol·min^−1^·mg^−1^ of C*b*_5_R. (Figure 1b). The dependence of the NADH:CoQ_0_ reductase activity with the concentration of CoQ_0_ was also examined using a fixed concentration of C*b*_5_R (1 µg/mL) and increasing concentration of CoQ_0_ (up to 2 mM) following the absorbance at 410 nm, in the presence of NADH (0.5 mM) and C*b*_5_R 1 µg/mL (Figure 1c). The dependence of the initial rate on the concentration of CoQ_0_ fitted to Michaelis-Menten model R^2^= 0.97 yielding a *V*_max_ value of 23.2 ± 2.1 µM/min and a *K_M_* value for CoQ_0_ of 292.8 ± 68.3 µM (Figure 1d).

**Figure 1.**
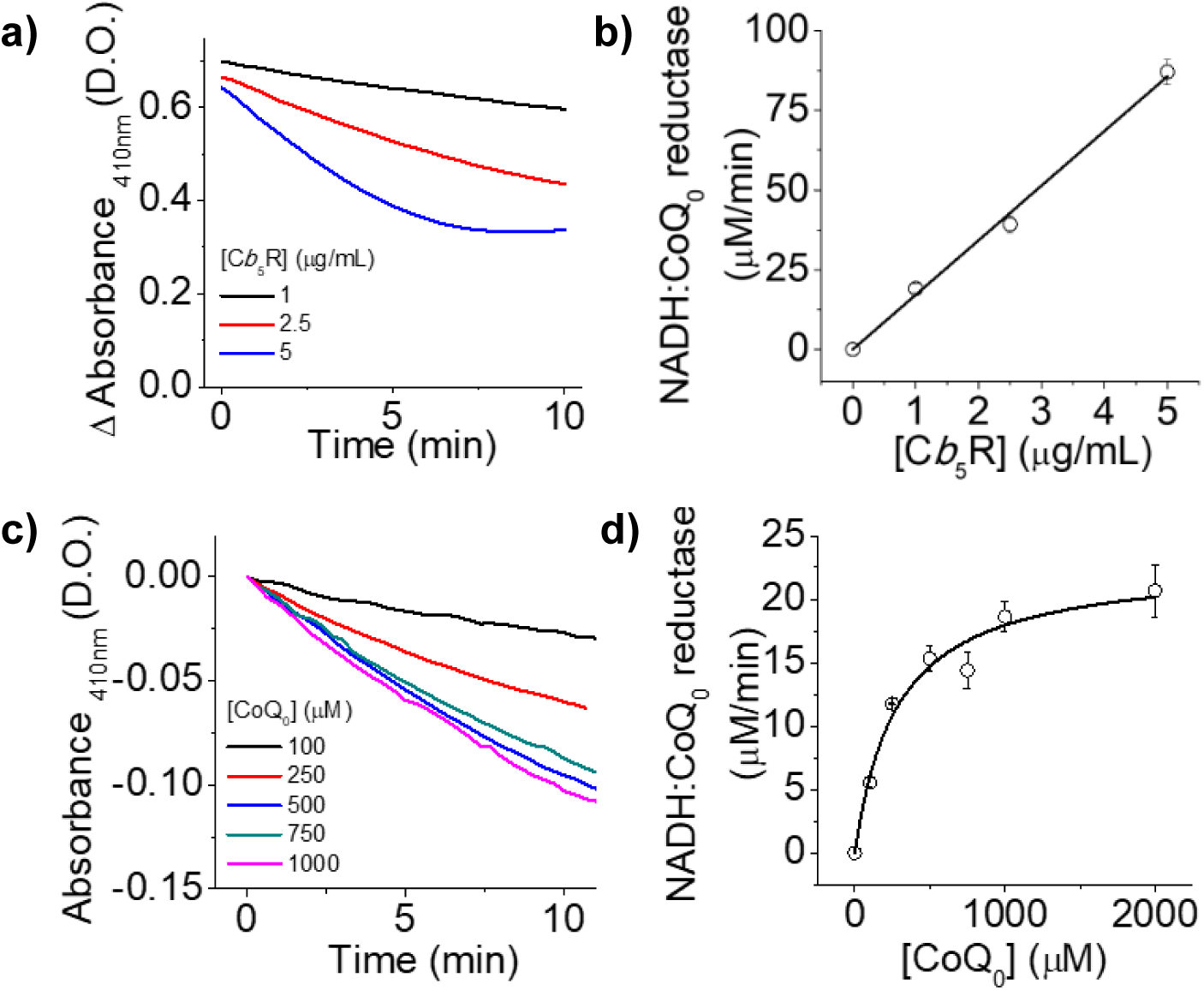
Dependence of NADH:CoQ_0_ reductase activity of C*b*_5_R upon enzyme and CoQ_0_ concentration. Panel a) shows absorbance values recorded at 410 nm as a function of time for different concentrations of C*b*_5_R: 1 (black line), 2.5 (red line), and 5 (blue line) µg/mL, at a fixed CoQ_0_ concentration (1 mM). Absorbance measurements were started after the addition of 500 µM NADH. Panel b) shows the maximum observed rate of CoQ_0_ reduction (µM/min) corresponding to the previously mentioned C*b*_5_R concentrations: 1, 2.5, and 5 µg/mL. The dotted line represents the best fit to a linear regression model. Panel c) shows the change in absorbance values at 410 nm relative to the initial recorded value, corresponding to 100, 250, 500, 750, 1000 µM CoQ_0_ concentrations, as a function of time. Panel d) displays the maximum CoQ_0_ reduction rate for the CoQ_0_ concentration used in panel c) at a fixed C*b*_5_R concentration (1 µg/mL). The dotted line represents the best fit to the Michaelis-Menten equation: *V*_0_ = (*V_max_* [CoQ_0_])/ (*K*_*m*_ + [CoQ_0_]).

### Molecular docking of C*b*_5_R complex with CoQ_0_

To further analyze the interaction between CoQ_0_ and C*b*_5_R in our experiments, molecular docking simulations were performed using Autodock Vina and visualized with UCSF Chimera. After performing multiple experiments (Supp. Table 1), the predicted binding energy corresponds to −5.8 kcal/mol. The predicted CoQ_0_ binding site in C*b*_5_R was identified by selecting all enzymes residues located within 5Å of the docked CoQ_0_ molecule. The residues obtained are: K110, Y112, G179, G180, T181, G182, C273, G274, P275, P276 and the FAD group where H-bonds are favored between C*b*_5_R amino acid residues and CoQ_0_ atoms: G180-O1, T181-O2, Y112-O1 and FAD-O3B-O1 (in H-bonds pairs residues from C*b*_5_R are labeled before the hyphen and CoQ_0_ atoms after) (Figure 2 and Supp. Table 1).

**Figure 2.**
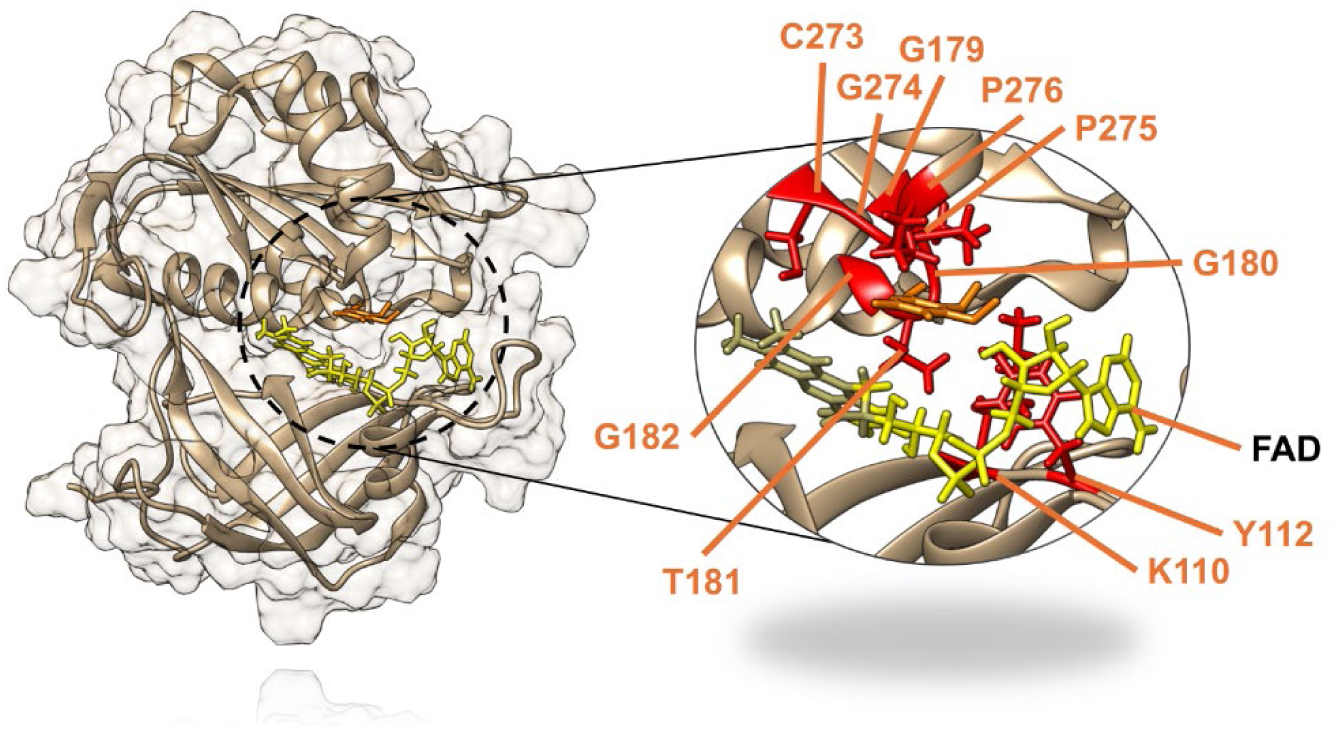
Molecular docking analysis of the CoQ_0_:C*b*_5_R complex. C*b*_5_R backbone is represented in brown, FAD group in yellow and CoQ_0_ in orange. The zoomed-in area on the right side of the panel shows the CoQ_0_ binding site in C*b*_5_R, where the side chains of the constituting amino acid residues are highlighted in red and labelled in orange with respect to the FAD group.

### Effect of CoQ_0_ on the NADH:C*b*_5_ reductase activity of C*b*_5_R

The NADH:C*b*_5_ reductase activity of C*b*_5_R was monitored by measuring the absorbance increase at 557nm in the presence of C*b*_5_ (30 µM), NADH (150 µM) and C*b*_5_R (0.05 µg/mL) (Figure 3a), as previously described in the Material and Methods section. The addition of CoQ_0_ was correlated with the decrease in the reduction rate and the concentration of reduced C*b*_5_. The presence of a lag phase, dependent on the concentration of CoQ_0_, and compatible with hysteretic behavior was observed (Figure 3 and 3c). The dependence of the lag phase duration (*τ*) on the concentration of CoQ_0_ fitted a linear regression model with a R^2^ = 0.97, and a slope of 1.51 ± 0.08 min per µM of CoQ_0_ (Figure 3b). Plotting the C*b*_5_ reduction rate against the concentration of CoQ_0_ yielded an IC_50_ value of CoQ_0_ of 1.7 ± 0.7 µM for the inhibition of this activity (Figure 3c) obtained with fixed concentrations of C*b*_5_ (30µM) and C*b*_5_R (0.05 µg/mL).

**Figure 3.**
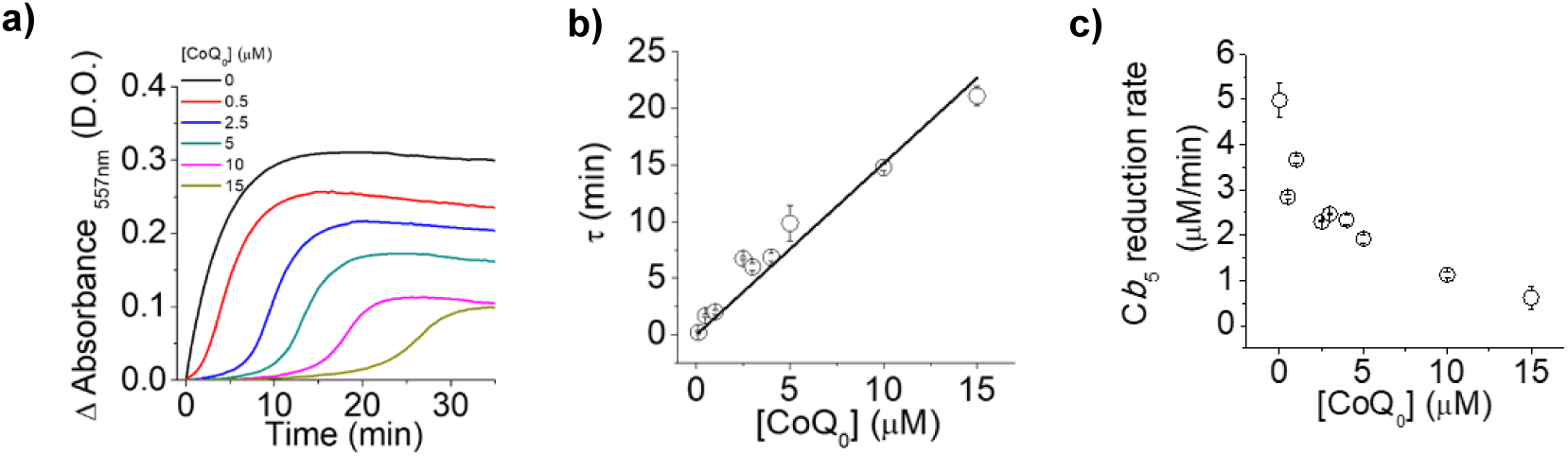
Dependence of NADH:C*b*_5_ reductase activity of C*b*_5_R on CoQ_0_ concentration. **Panel (a)** shows kinetic of C*b*_5_ reduction measured by the absorbance increment recorded at 557 nm over time in the presence of different CoQ_0_ concentrations: 0 (black line), 0.5 (red line), 2.5 (blue line), 5 (green line), 10 (pink line), and 15µM (yellow line), at fixed concentrations of C*b*_5_ (30 µM) and C*b*_5_R (50 ng/mL). Measurements in all other experimental conditions started after the addition of 150 µM NADH to the assay. **Panel (b)** displays the duration (in minutes) of the lag phase (*τ*) as a function of the added CoQ_0_ concentration. The dotted line represents the best fit to a linear regression model. **Panel (c)** shows the maximum observed rate of C*b*_5_ reduction (µM/min) as a function of the CoQ_0_ concentration present in the assay.

### Effect of C*b*_5_R concentration on the modulated NADH:C*b*_5_ reductase activity by CoQ_0_

The reduction of C*b*_5_ using different concentrations of C*b*_5_R (0.005-0.050 µg/mL) and fixed concentration of C*b*_5_ and NADH followed a pseudo-first order kinetic (Figure 4a). In the presence of CoQ_0_ (5 µM), increasing the concentration of C*b*_5_R reversed the hysteretic effect by increasing the rate of C*b*_5_ reduction and a decrease in the lag phase (Figure 4a and b). The dependence of *τ* with the concentration of C*b*_5_R fitted to an exponential decay of form: 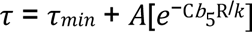 with a R^2^ of 0.98 (Figure 4b) that allowed us to determine a theorical *τ_min_* of 10.7 ± 4.9min, a *k* value of 0.009 ± 0.004 µg/mL and a decay amplitude (A) of 73.9 ±28.7 min. In the absence and presence of CoQ_0_, a linear dependence of the activity on the concentration of enzyme was observed (R^2^ = 0.99 and 0.86, respectively) with slopes of 97.4± 2.5 and 28.4 ± 4.1 µmol/min/mg, respectively (Figure 4c), correlating with a 70.8% decrease in the activity induced by CoQ_0_ (5µM) in the dependence of the C*b*_5_ reduction rate with the concentration of C*b*_5_R.

**Figure 4.**
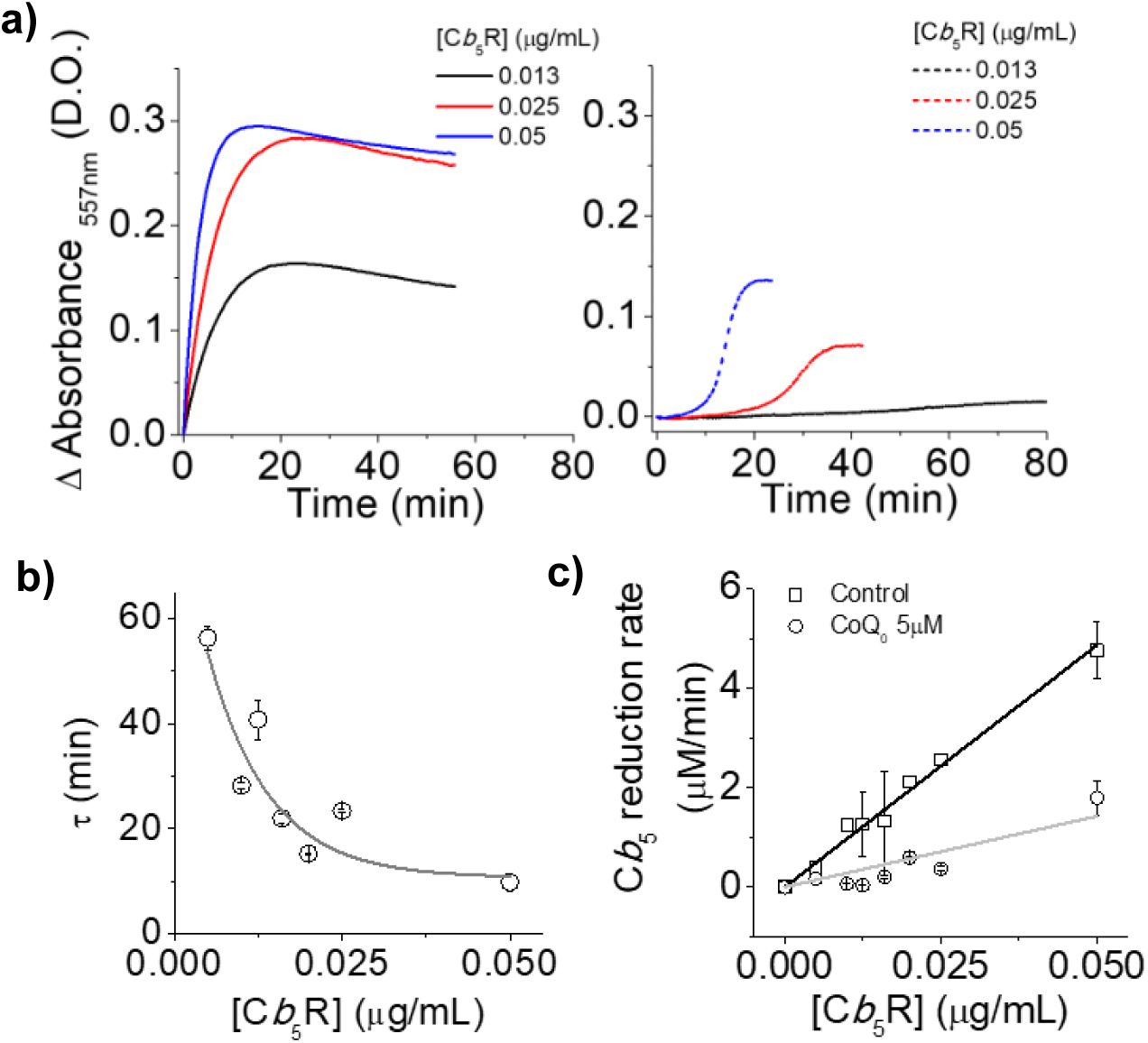
Kinetics of the dependence of C*b*_5_ reduction on C*b*_5_R concentration in the absence and presence of CoQ_0_. **Panel a),** on the left, shows the increment of C*b*_5_ absorbance at 557 nm as a function of time in the absence (left side) of CoQ_0_ at different C*b*_5_R concentrations: 0.013 (black line), 0.025 (red line) and 0.05 µg/mL (blue line) measured with a fixed concentration of C*b*_5_ (30 µM). Absorbance measurements started by the addition of 150 µM NADH to the assay. **Panel a),** on the right, shows the dependence of C*b*_5_ reduction rate on the same C*b*_5_R concentration in the presence of CoQ_0_ (5 µM)**. Panel b** represents the duration (in minutes) of the lag phase (*τ*) recorded at different C*b*_5_R concentrations after the addition of 5 µM CoQ_0_. The line represents the best fit to the exponential decay equation: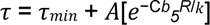. **Panel c** represents the dependence of the C*b*_5_ reduction activity on C*b*_5_R concentrations in the absence (open squares) and presence of 5 µM CoQ_0_ (open circles). The lines represent the best fits to the linear regression equation.

### Effect of C*b*_5_ concentration on the modulated NADH:C*b*_5_ reductase activity by CoQ_0_

The dependence of the NADH:C*b*_5_ reductase activity of C*b*_5_R upon C*b*_5_ concentration (from 5-35 µM) using different concentrations of CoQ_0_ (0, 1 and 5 µM) and a fixed concentration of NADH (150 µM) and C*b*_5_R (0.05 µg/mL) is shown in Figure 5. The kinetics of the dependence of the activity on C*b*_5_ in the absence of CoQ_0_ followed a pseudo-first order saturation kinetics (Figure 5a) while in the presence of CoQ_0_ (1 and 5 µM) hysteretic behavior was observed (Figure 5b), characterized by a lag phase in the reduction of C*b*_5_ with the concentration of CoQ_0_, that was more prominent at lower C*b*_5_ concentrations and diminished at higher C*b*_5_ concentrations.

**Figure 5.**
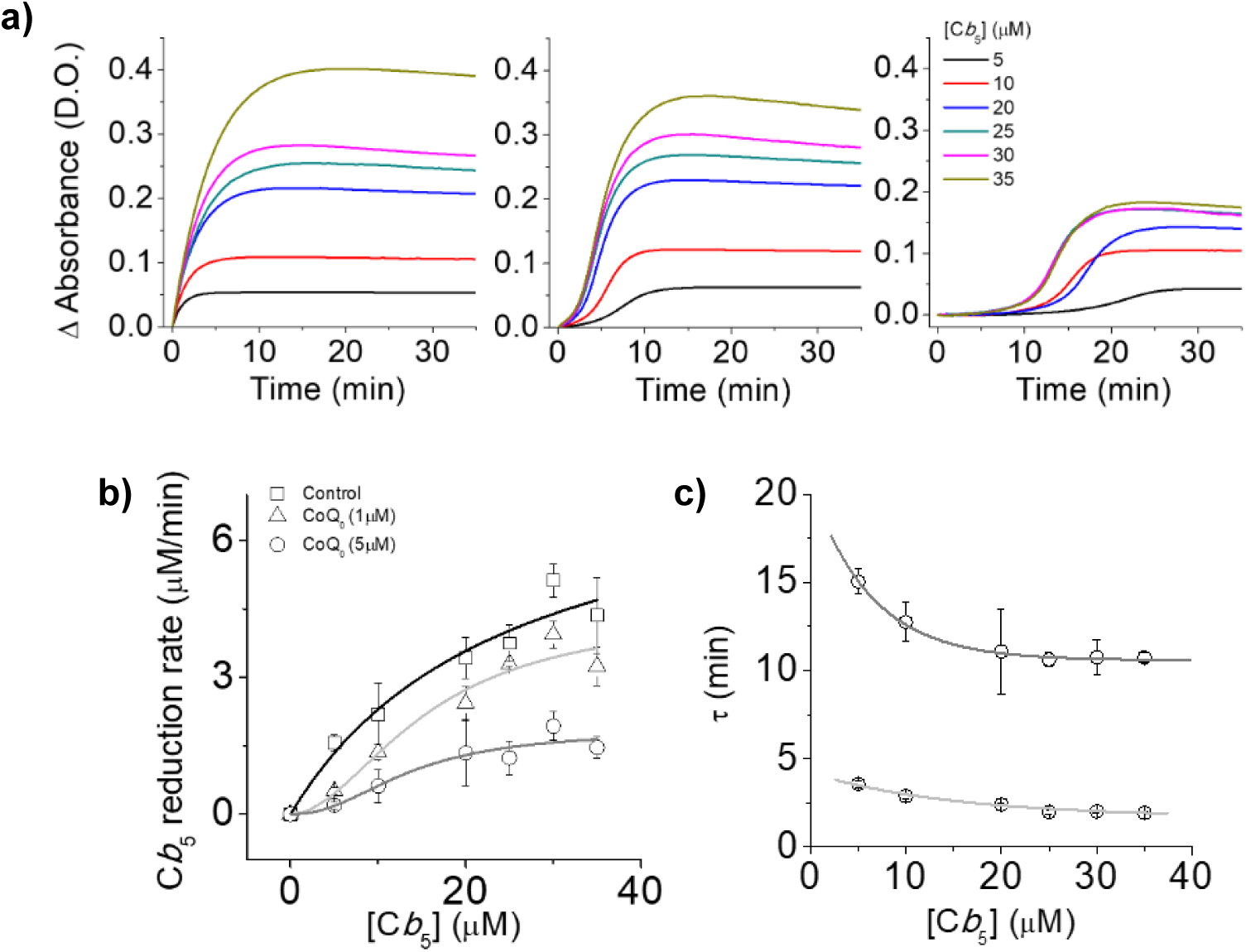
Dependence of C*b*_5_ reduction kinetics by C*b*_5_R on C*b*_5_ concentration in the absence and presence of CoQ_0_. **Panels a)** show absorbance values recorded at 557 nm as a function of time, for three CoQ_0_ concentrations: 0 µM (left side), 1 µM (middle side), and 5 µM (right side); at different C*b*_5_ concentrations: 5 (red line), 10 (orange line), 20 (dark green line), 25 (light green line), 30 (light blue line), 35 (dark blue line), and 40 µM (purple line); and at a fixed C*b*_5_R concentration (50 ng/mL). Absorbance measurements began after the addition of 150 µM NADH. **Panel b)** represents the maximum Cb_5_ reduction rate plotted as a function of the C*b*_5_ concentration in assays in the presence of different CoQ_0_ concentrations: 0 (open squares), 1 (open triangles), and 5 µM (open circles). In this panel, the grey lines represent the best fit to the Hill equation: *V*_₀_ = (*V*_max_ [C*b*_5_])^n^ / (*K*_ₘ_ + [C*b*_5_])^n^. **Panel c)** show the duration of the lag phase (*τ*) recorded as a function of C*b*_5_ concentration after the addition of 1 µM (light grey) and 5 µM (dark gray) CoQ_0_. In this panel, lines represent the best fit to the exponential decay equation: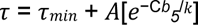.

We plotted the dependence of the *τ* value on the C*b*_5_ concentration in the presence of CoQ_0_ (1 and 5 µM) (Figure 5c). For both CoQ_0_ concentrations, the data showed a decrease in the *τ* value with increasing C*b*_5_ concentration, although this dependence did not fit well to a linear regression, hyperbolic or an exponential decay model, as previously observed for the dependence of the activity on the concentration of enzyme (Figure 4c). The dependence of *τ* on the concentration of C*b*_5_ fitted to an exponential decay model with a theorical *τ*_min_ of 1.7 ± 0.2 min, a *k* value of 15.2 ± 4.2 µM and a A value of 2.5 ± 0.3 min for assays performed with CoQ_0_ 1µM. In the presence of CoQ_0_ (5 µM), we determined a *τ*_min_ of 10.6 ± 0.1min, a *k* value of 6.2 ± 1.2 µM and an A value of 9.9 ±1.7 min.

The dependence of the C*b*_5_ reduction rate (Figure 5b, open squares) on the concentration of C*b*_5_, in the absence of CoQ_0_, fitted the Hill equation in the absence and presence of CoQ_0_ (1 and 5 µM, open triangles and circles, respectively) R^2^ of 0.98, 0.97 and 0.96, respectively.

In the absence of CoQ_0_, we obtained a *V*_max_ value of 8.9 ± 1.1 µM/min and a *K*_M_ value for C*b*_5_ of 25.0 ± 1.1 µM and a cooperativity index (*n* value) of 1.0 ± 0.1. In the presence of 1 µM of CoQ_0_, the *V*_max_ value was 4.6 ± 0.8 µM/min and the *K*_M_ value for C*b*_5_ was 14.9 ± 3.7 µM. We determined an *n* value of 1.8 ± 0.4. Similar values were obtained with 5 µM of CoQ_0_ (*V*_max_ = 1.8 ± 0.2; *K*_M_ =10.9 ± 2.1 and a *n* value of 2.0 ± 0.5) (Table 1).

**Table 1.**
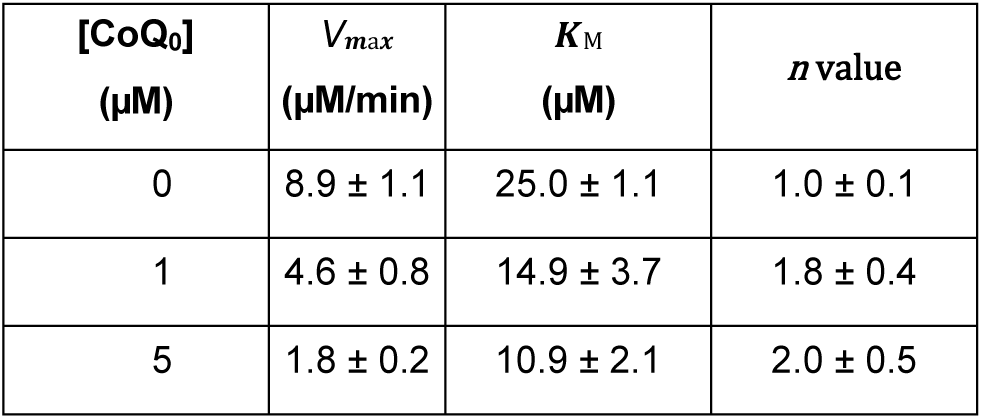
Calculated values for the parameters: *V*_max_ (maximum velocity), *K*_M_ (Michaelis-Menten constant), cooperativity coefficient. For parameters calculation the Hill equation was used: y=(*V*_max_[C*b*_5_])^n^/(*K*_M_+[C*b*_5_])^n^.

| [CoQ<sub>0</sub>]<br>($\mu\text{M}$) | $V_{\max}$<br>($\mu\text{M}/\text{min}$) | $K_M$<br>($\mu\text{M}$) | $n$ value |
| --- | --- | --- | --- |
| 0 | $8.9 \pm 1.1$ | $25.0 \pm 1.1$ | $1.0 \pm 0.1$ |
| 1 | $4.6 \pm 0.8$ | $14.9 \pm 3.7$ | $1.8 \pm 0.4$ |
| 5 | $1.8 \pm 0.2$ | $10.9 \pm 2.1$ | $2.0 \pm 0.5$ |

### Measurement of the kinetic of NADH consumption in the presence of C*b*_5_ and CoQ_0_

We evaluated the kinetics of NADH consumption by the presence of NADH (50 µM) in the absence or presence of C*b*_5_ (30 µM), CoQ_0_ (5 µM) and a fixed concentration of C*b*_5_R (0.1 µg/mL) (Figure 6). In the absence of CoQ_0_ and C*b*_5_, the oxidation of NADH was near the limit of detection of the used methodology but initial oxidation rates were estimated at 0.17 ± 0.05 µM/min and 0.28 ± 0.04 µM/min, respectively. The addition of C*b*_5_ (30 µM) increased the initial rates of NADH oxidation to 3.8 ± 0.3 µM/min and in the presence of CoQ_0_ dropped to 2.4 ± 1.1 µM/min.

**Figure 6.**
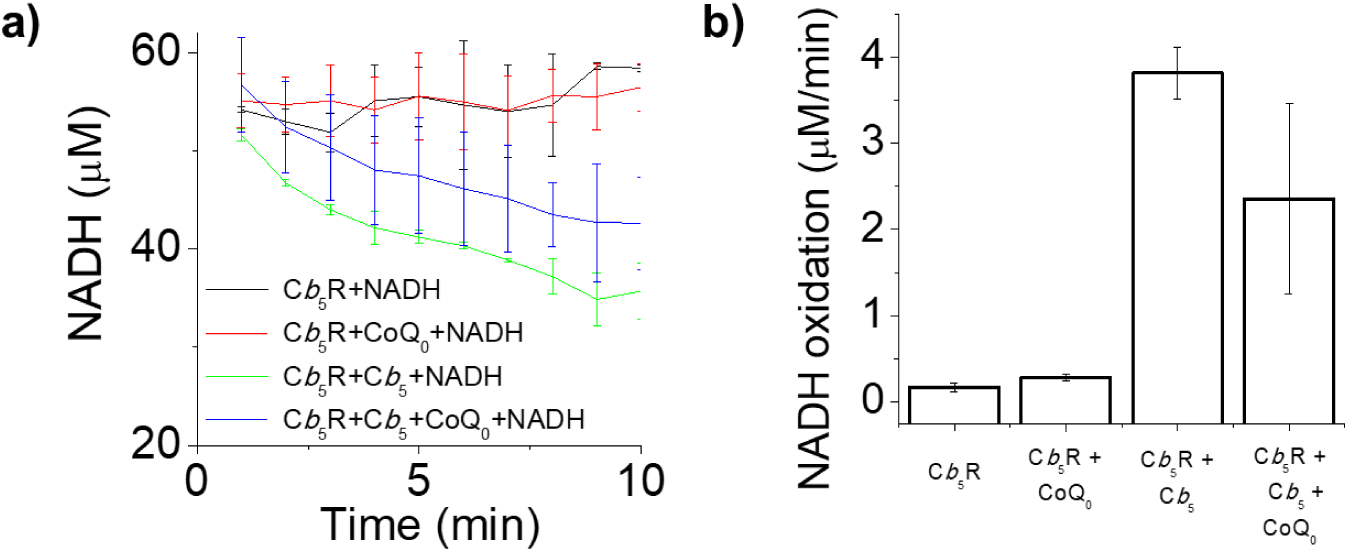
Dependence of NADH consumption on C*b*_5_R activities and the influence of CoQ_0_ on the assays. **Panel a)** shows the kinetic of NADH consumption after addition of NADH to the assay using a C*b*_5_R concentration of 0.1 µg/mL in the absence (black line) and presence of 30µM of C*b*_5_ (green line) and 5µM of CoQ_0_ (red line) or both substrates (blue line). **Panel b)** displays the maximum NADH oxidation rate (µM/min) obtained under the different experimental conditions described in panel a).

### Electrochemistry

Cyclic voltammetry (CV) experiments were designed to individually characterize the proteins and quinone redox potentials, enabling later comparison of their properties in mixtures. The components were analyzed within the 0 V to 0.6 V window, returning to −0.7 (*vs* NHE), at different scan rates. The independent voltammograms enabled us to calculate the formal reduction potentials (E^0^’) for C*b*_5_ and CoQ_0_ (Figure 7, panels A and B, respectively).

**Figure 7.**
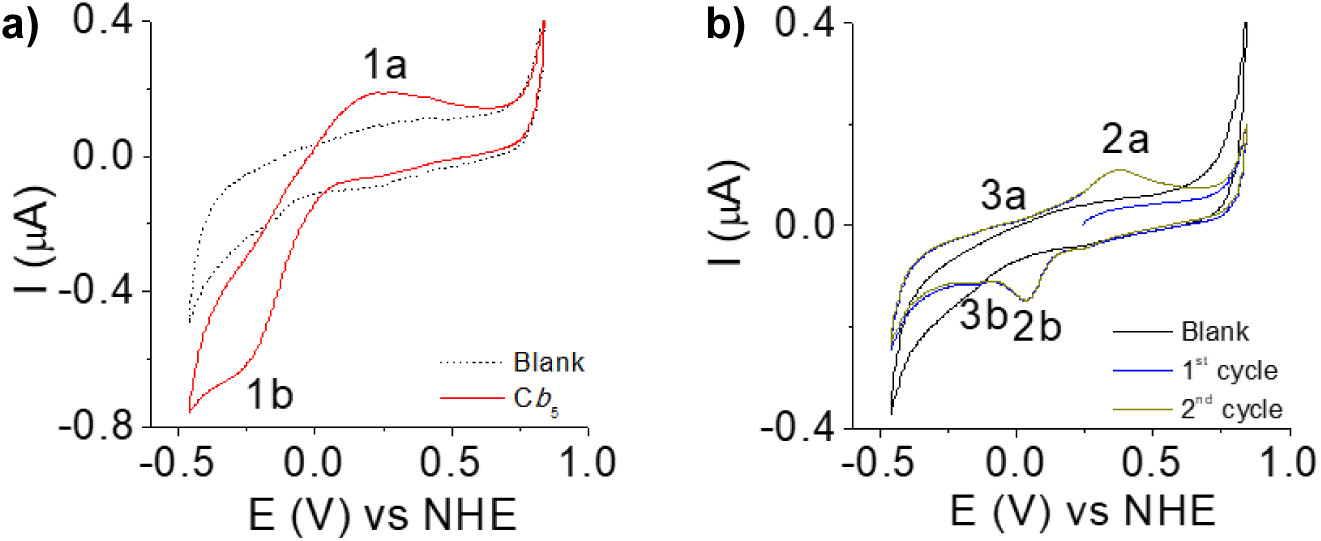
Cyclic voltammetry of C*b*_5_ and CoQ_0_. **Panel a)** C*b*_5_’s representative cyclic voltammetry (50 mV/s). 2^nd^ cycle voltammograms (Blank and C*b*_5_), of the assays with C*b*_5_ (150 µM) entrapped in a 3.5 kDa cellulose membrane with 2 mM of neomycin in 20 mM phosphate buffer, 100 mM of NaCl and 1 mM of EDTA as electrolyte. **Panel b**) CoQ_0_’s cyclic voltammetry (20 mV/s). Comparison between 1^st^ and 2^nd^ voltammograms of the assays with CoQ_0_ (30 µM) in the presence of a 3.5 kDa cellulose membrane with 2 mM of neomycin and 20 mM phosphate buffer, 100 mM of NaCl and 1 mM of EDTA as electrolyte with the control (blank) CV.

A E^0’^ value of −13 ± 23 mV vs NHE at pH 7 was obtained from voltammograms of C*b*_5_ (Figure 7a). When analyzing CoQ_0_ in 1^st^ and 2^nd^ voltammograms cycles, no oxidation peak was observed. In the 2^nd^ cycle, two processes were observed (denoted as 2 and 3; process 3 presenting less current intensity). Contrary to the 1^st^ cycle, in the second cycle, an oxidation peak (2a) was observed, and associated with the previous reduction of the CoQ_0_ in the 1^st^ cycle (in the cathodic direction, from 0.841 to −0.459 V vs NHE), implying that the existence of a larger population of reduced CoQ_0_, is then present to be oxidized in the 2^nd^ cycle (Figure 7b). From the voltammograms, it was possible to obtain for process 2 the formal potential, E^0’^, of +207 ± 7 mV vs NHE at pH 7. For C*b*_5_R, the data did not allow us to determine its redox potential parameters (data not shown).

To better understand the mechanism observed in this work, mixtures of various components were analyzed. The data obtained from the complete mixtures of C*b*_5_R + C*b*_5_ + CoQ_0_ revealed a decrease of the C*b*_5_ cathodic peak intensity (Figure 8a, red) when in the presence of C*b*_5_R (green line), suggesting a decrease in oxidized C*b*_5_ molecules in solution, as a result of electron transfer from C*b*_5_R to C*b*_5_. This observation may also be due to additional components in the mixture, implying that less electrode surface is available to react with oxygen. In contrast, the addition of CoQ_0_ over the C*b*_5_R and C*b*_5_ mixture, there is again a slight increase in the Cb5 cathodic current (Figure 8a, brown line Figure 8b). The interaction between CoQ_0_ with the C*b*_5_/C*b*_5_R system was also analyzed across the 1^st^ and 2^nd^ CV cycles (Figure 8 c and d). The quinone was observed to be slightly more oxidized (1^st^ cycles, blue line) in the presence of C*b*_5_R than when tested alone (red line). This was highlighted by the absence of anodic processes in the 1^st^ anodic scan (from 0.241 to 0.841 V vs NHE) in the CoQ_0_+C*b*_5_R mixtures, suggesting that CoQ_0_ interacts with C*b*_5_R by donating electrons to the enzyme. Notably, the C*b*_5_R + CoQ_0_ mixture alone generates a 3^rd^ oxidation signal (4a, blue, better observed in the 2^nd^ cycles) that was not visible in the other mixtures. This peak may indicate complex formation between the two molecules, or the formation of new species resulting from their interaction. When compared to the complete mixture (C*b*_5_R + C*b*_5_ + CoQ_0_), the current waves corresponding to the quinone processes exhibit a larger potential separation (increase ΔE_p_) in the presence of C*b*_5_ compared to C*b*_5_R alone, which might suggest that interaction between the two could alter the redox potentials. The addition of C*b*_5_ to the mixture seems to lower the quinone’s oxidation peak, which may be an indication that the enzyme prefers to interact with C*b*_5_ over CoQ_0_, as an electron acceptor.

**Figure 8.**
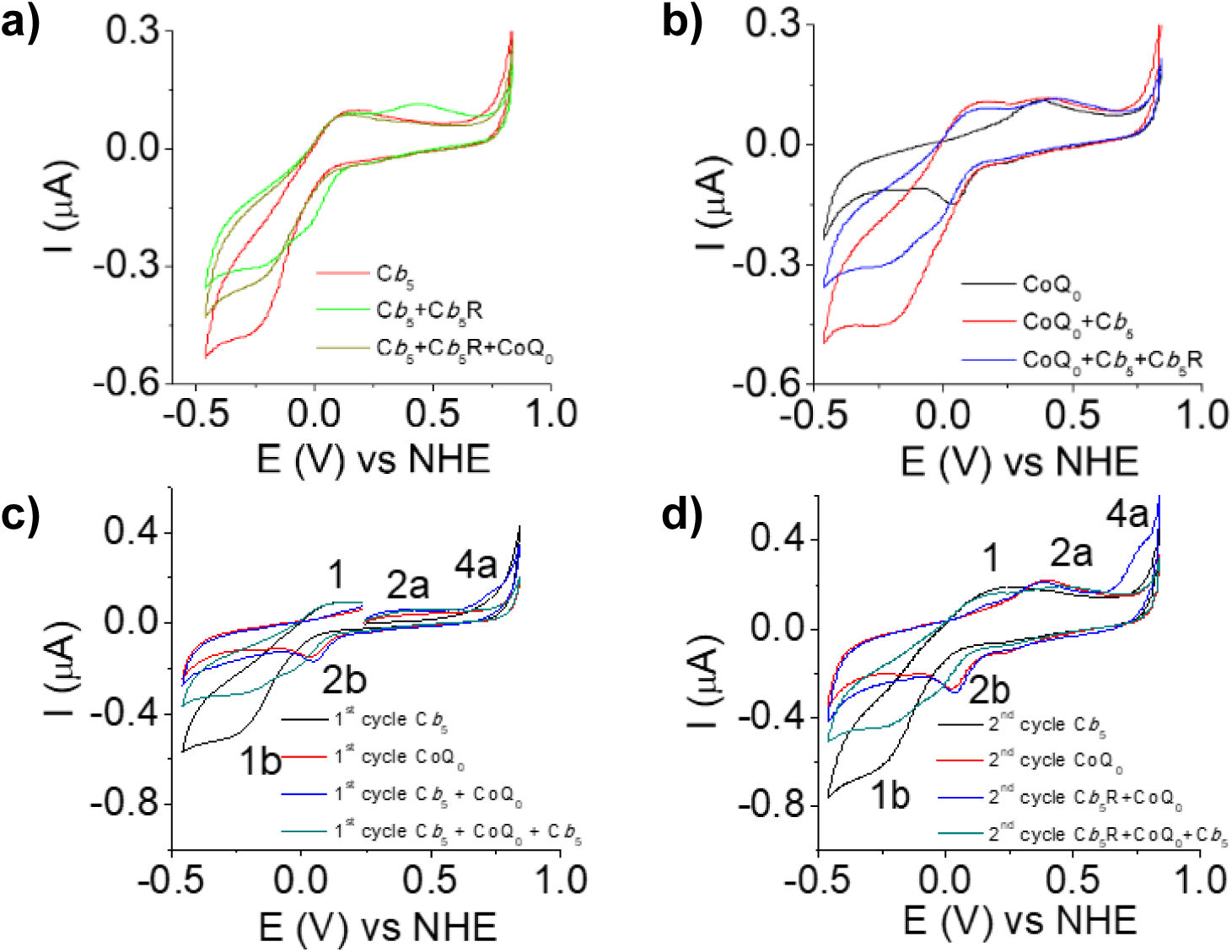
Cyclic voltammetry of mixtures of compound’s mixtures studied in this work, in the presence of C*b*_5_R. **Panel a)** Comparison between the 2^nd^ cycle of the voltammogram of C*b*_5_, C*b*_5_R + C*b*_5_ and C*b*_5_R + CoQ_0_+ C*b*_5_ at 20 mV/s. **Panel b)** Comparison between the 2^nd^ cycle voltammogram of CoQ_0_, CoQ_0_+ C*b*_5_ and C*b*_5_R + CoQ_0_+ C*b*_5_ at 20 mV/s. *Panel c) 1^st^ cycles and* Panel d) 2^nd^ cycles of representative voltammograms of the samples: C*b*_5_, CoQ_0_, C*b*_5_R + C*b*_5_ and C*b*_5_R + CoQ_0_+ C*b*_5_ at 20 mV/s. All scans started in the anodic direction.

## DISCUSSION

C*b*_5_R is referred as a pleiotropic enzyme since it catalyzes multiple reduction reactions [8,13]. In this work, we confirmed that C*b*_5_R has an NADH:CoQ reductase activity, consistent with studies suggesting a role in the regulation of cellular oxidative stress through the reduction of CoQ_10_ [19,35]. The linear dependence of the NADH:CoQ_0_ reductase activity on C*b*_5_R concentration and the relationship between the reduction rate and the concentration of CoQ_0_ allowed us to obtain the kinetic parameters via fitting to the Michaelin-Menten equation. The obtained *V*_max_ value of 23.2 ± 2.1 µM/min and *K_M_* value for CoQ_0_ of 291.8 ± 68.3 µM, indicate low affinity of C*b*_5_R for CoQ_0_ compared to C*b*_5_ (*K_M_* value: 12.8 ± 2.3 µM). Reported CoQ_10_ levels in tissues (i.e.: 0.1-2 nmol/mg of protein [36] and 1-2 µM in plasma [37]) and cells (i.e.: approx. 150 ng/mg of protein in keratinocytes [38]) can increase after supplementation [39], and are comparable to those reported for C*b*_5_: 0.27 nmol/mg protein [40] and 0.26 µM in human liver microsomes and human red cells [41], respectively. Therefore, the calculated *K_M_* values for CoQ_0_ and C*b*_5_ suggest that their reduction by the C*b*_5_R using NADH will not operate at *V*_max_ in biological systems, since both C*b*_5_ and CoQ_10_ concentrations, reported in cells and tissues, are below those used in this study. Nevertheless, the activity might be modulated by small changes on CoQ_10_ concentration *in vivo*. Moreover, comparison of the *K*_M_ values for C*b*_5_ and CoQ_0_ suggests that in model systems where substrates are tested independently in the assays, the enzyme will preferentially reduce C*b*_5_ over CoQ_0_. The docking data support a binding site for CoQ_0_ near the catalytic site: K110, Y112, G179, G180, T181, G182, C273, G274, P275, P276 and the FAD group, featuring hydrogen bonding with Y112, G180, T181 and the FAD group. This binding site is similar to that reported for some inhibitors of C*b*_5_R that we have characterized [25,42].

Despite the protective role attributed to C*b*_5_R through the NADH:CoQ reductase activity, we previously reported that some quinones can uncouple the NADH:C*b*_5_ reductase activity of C*b*_5_R [43]. On these grounds, it was relevant to know whether ubiquinone, one of the main biological quinones in tissues, and CoQ_0_ as its soluble derivate, could affect this activity. Moreover, one of the objectives of this work was to assess whether CoQ_0_ could hysteretically modulate the NADH:C*b*_5_ reductase activity of C*b*_5_R, based on previous findings showing that the NADH-dependent reduction of C*b*_5_ in microsomes, where several enzymes may contribute to this activity, is hysteretically modulated by CoQ_0_ and CoQ_10_[22]. In this work, we measured this activity using the recombinant human soluble C*b*_5_R and C*b*_5_, to avoid artifacts associated with mixing soluble and membrane isolated components and using stoichiometric ratios of C*b*_5_ to quinone in the 1:1, 1:10 range, compatible with their reported levels in biological samples [36,37,40,41].

We first characterized the NADH:C*b*_5_ reductase activity of C*b*_5_R in the presence of different concentrations of C*b*_5_R, C*b*_5_ and CoQ_0_. In the absence of CoQ_0_, the reduction of C*b*_5_ follows pseudo-first-order kinetics consistent with bimolecular reaction requiring both NADH and C*b*_5_, where the concentration of NADH is in excess (150 µM) compared to C*b*_5_. Under these conditions, the C*b*_5_ reduction rate depends exclusively on the C*b*_5_ concentration present in the assays which simplifies the kinetic model. The appearance of a lag phase (*τ*) in the C*b*_5_ reduction kinetic upon the addition of CoQ_0_, is compatible with a previously undescribed role of CoQ_0_ as a hysteretic modulator of the NADH:C*b*_5_ reductase activity of C*b*_5_R. The duration of *τ* exhibits a linear dependence on the concentration of CoQ_0_ (increasing) and decreases with increasing concentrations of C*b*_5_. Increasing C*b*_5_R concentration induced a decrease in the *τ* value fitting an exponential decay equation.

It should be noted that the rate of C*b*_5_ reduction by C*b*_5_R during the lag phase was always lower than its maximum rates observed at the stationary phase of the kinetic. This behavior indicated that the slow transition induced by CoQ_0_ involves at least one inactive or partially active catalytic state transitioning to the state [44]. Regarding the effect of CoQ_0_ on the maximum C*b*_5_ reduction rate, the kinetic parameters decreased compared to conditions without CoQ_0_, as demonstrated in the assays assessing these parameters dependence on concentrations of CoQ_0_, C*b*_5_R and C*b*_5_.

The effect of CoQ_0_ addition on the maximum C*b*_5_ reduction rate was also studied. Parameters showed a decrease when the dependence of the NADH: C*b*_5_ reductase was measured varying with CoQ_0_, C*b*_5_R and C*b*_5_ concentrations relative to the control (no CoQ_0_ added). This evidence demonstrates that the modulation by CoQ_0_ results from the inhibition of the NADH:C*b*_5_ reductase activity of the C*b*_5_R. The analysis of the dependence of the C*b*_5_ reduction rate on CoQ_0_ and C*b*_5_ concentrations revealed substrate concentration-dependent kinetics. The maximum rate of C*b*_5_ reduction exhibited a linear dependence on C*b*_5_R concentration, and data fitting yielded a drop of 70.8% in catalytic activity upon addition of 5 µM of CoQ_0_. Additionally, we evaluated CoQ_0_ ability to inhibit C*b*_5_R’s NADH: C*b*_5_ reductase activity of C*b*_5_R by calculating the IC_50_ value associated with the decrease in the C*b*_5_ reduction rate using 30 µM of C*b*_5_ in the assays, which was 1.7 ± 0.7 µM of CoQ_0_.

The reduction rate of C*b*_5_ by C*b_5_*R as a function of C*b_5_* concentration in the absence and presence of CoQ_0_ (1 and 5 µM) was fitted to the Hill equation yielding kinetic parameters for our assays (Table 1). The analysis of the *V*_max_ values revealed and inhibition of the reduction rate that was dependent on the concentration of CoQ_0_. Our data also revealed that at higher CoQ_0_ concentrations, *K*_M_ values decreased, accompanied by cooperativity for C*b*_5_ (*n* values near 2 with CoQ_0_ 1 and 5 µM) supporting increased affinity for C*b*_5_, due to formation of a potential CoQ_0_:C*b*_5_R complex, altering the enzyme binding. To shed light on this kinetic behavior, we performed electrochemical assays (cyclic voltammetry). The cyclic voltammograms of C*b*_5_ enabled us to calculate an E0’ value of −13 ± 23 mV vs NHE at pH 7. For CoQ_0_, an E^0’^ of +207 ± 7 mV vs NHE at pH 7 was obtained for process 2. For this process it was possible to calculated the formal potential, from 2a and 2b current peaks, that were assessed to correspond the semiubiquinone state [45]. For process 3, namely considering peaks 3a and 3b (confirmed to be a redox pair), a formal potential of approximately −156 mV vs NHE were calculated, which closely match the literature result for the CoQ/CoQH_2_ redox pair [46]. For C*b*_5_R, it was not possible to determine parameters as done for the other two components (C*b*_5_ and CoQ_0_) and using other type of electrodes [47]. In the literature, however, some assays with this enzyme were performed on thiol modified gold electrodes [47]. In our results, with pyrolytic graphite, the voltammogram present a large cathodic current observed around −0.449 V vs NHE when C*b*_5_R is present. This may occur due to an interaction between C*b*_5_R and produced oxygen species b the electrode or by the C*b*_5_R itself [42,48]. When compared to the complete C*b*_5_R + C*b*_5_ + CoQ_0_ mixture, the peaks corresponding to quinones processes exhibit greater separation reflecting a possible interaction between C*b*_5_ and CoQ_0_ that alters the redox potentials The addition of C*b*_5_ to the mixture also lowers the quinone’s oxidation peak. This decrease in quinone oxidation indicates that the enzyme prefers C*b*_5_ over CoQ_0_, as an electron acceptor and may support the formation of alternative CoQ_0_/protein complexes, potentially responsible for the hysteretic behavior. Further experiments are needed to fully elucidate the mechanism.

Finally, NADH consumption data confirm that both CoQ_0_ and C*b*_5_ increase the oxidation of NADH, as a result of the presence of an electron acceptor different to oxygen, correlating with the concentration of the acceptor used in each assay (5µM of CoQ_0_ and 30µM of C*b*_5_). Noteworthy, the presence of CoQ_0_ and C*b*_5_ in the assays, implies a slight decrease of the NADH oxidation rate that was measured relative to C*b*_5_ alone, suggesting a deceleration of the catalytic cycle due to the hysteretic modulation induced by CoQ_0_ potentially related to the full reduction of the flavoprotein via the flavin semiquinone reduction.

## Conclusion

Overall, this work demonstrates that CoQ_0_ acts as a hysteretic modulator of the NADH:C*b*_5_ reductase activity of C*b*_5_R, through probably an alternative conformation induced on C*b*_5_R, leading to an alternative CoQ_0_-dependent one electron reduction pathway, decelerating both the oxidation of NADH and C*b*_5_ reduction. Our results support the importance of measurements of endogenous components, including C*b*_5_R activity, C*b*_5_ and quinones levels on biological samples, to evaluate the outcomes in the pathways where C*b*_5_ participate. This might be key for redox control on metabolic pathways, since the reported modulation described in this manuscript may give important indicators for C*b*_5_-dependent pathways including fatty acids desaturation, xenobiotic detoxification and those associated with pathologies coursing through changes in the levels of C*b*_5_ and associated oxidative stress [49–53], severe systemic neurological dysfunction [50,54,55] and detoxification of xenobiotics [56,57].

## Data availability

The data that support the findings of this study are available from the corresponding author, AKSA, upon reasonable request.

## Acknowledgement

Open Access funding provided thanks to the CRUE-CSIC agreement with [ACS/ Elsevier / IEEE / RSC / Springer / Wiley]. GNV acknowledges FCT/MCTES for funding through grant number 2021.06216.BD. We thank Prof. Carlos Gutiérrez Merino from the Instituto de Biomarcadores de Patologías Moleculares, Universidad de Extremadura, for providing essential material for this work. This work (CMC and GNV) was financed by national funds from FCT - Fundação para a Ciência e a Tecnologia, I.P., under the scope of the project UID/50006/2025 of the Associate Laboratory for Green Chemistry - LAQV REQUIMTE.

