## Supplementary material for "Ubiquinone is a hysteretic modulator of the NADH:cytochrome *b*_5_ reductase activity of human C*b*_5_R": Supp. information

**Supplementary information**

| **Experiment 1** | | | | |
| --- | --- | --- | --- | --- |
| **Binding energy (kcal/mol)** | **Cluster** | **Pose** | **C*b*_5_R’s residues forming the binding site** | **H-bonds**  **C*b*_5_R-CoQ_0_** |
| **-5.8** | **1** | **1** | **FAD, K110, Y112, G179, G180, T181, G182, I183, C273, G274, P276** | **G180-O1** |
| **-5.7** |  | **2** | **FAD, Y112, G179, T181, G182, C273, G274, P275, P276,** F300 | **T181-O2** |
| **-5.3** |  | **4** | **FAD, K110, Y112, G179, G180, T181, G182, , C273, G274, P275, P276** | **FAD-O3B-O1** |
| **-4.8** |  | **6** | **FAD, K110, Y112, G179, G180, T181, G182, C273, G274, P275, P276** | **Y112-O1, FAD-O3B-O1** |
| **-4.6** |  | **9** | **FAD, K110, Y112, G179, G180,T181, G182, C273, G274, P275, P276**, F300 |  |
| -5.6 |  | 3 | **G179**, A208, N209, Q210, T237, G250, F251, V252, **P275**, P277, M278, Y281, A282 |  |
| -5.0 |  | 5 | **FAD**, H54, D55, **K110**, V111, **Y112**, F113, T116, Q210, D214 |  |
| -4.7 |  | 7 | **FAD**, H54, D55, **K110**, V111, **Y112**, F113, T116, Q210 |  |
| -4.7 |  | 8 | A208, **Q210**, D239, F251, V252, **P275**, P277, M278, Y281, A282 |  |
| **Experiment 2** | | | | |
| **Binding energy (kcal/mol)** | **Cluster** | **Pose** | **C*b*_5_R’s residues forming the binding site** | **H-bonds**  **C*b*_5_R-CoQ_0_** |
| **-5.8** | **1** | **1** | **FAD, K110, Y112, G179, G180, T181, G182, I183, C273, G274, P275, P276** |  |
| **-5.7** |  | **2** | **FAD. K110,Y112, G179, G180,T181, G182, C273, G274, P275, P276,**P277 | **T181-O2** |
| **-5.2** |  | **5** | **FAD,K110, Y112, G179, G180, T181, G182, C273, G274, P275, P276** |  |
| **-5.1** |  | **6** | **FAD,K110, Y112, G179, G180, T181, G182, C273, G274, P275, P276** |  |
| **-5.0** |  | **7** | **FAD, K110, Y112, G179, G180, T181, G182,** Q210, D214**,C273, G274, P275, P276** |  |
| **-5.6** |  | 3 | **G179**, A208, N209, Q210, T237, D239, G250, F251, V252, **P275**, P277, M278, Y281, A28 |  |
| **-5.2** |  | 4 | **G179, G180,** A208, N209, Q210, T237, L238, D239, F251, V252, **P275,** P277, M278, A282 |  |
| **-5.0** |  | 8 | **FAD,** H54, D55, **K110,** V111, **Y112,** F113, T116 |  |
| **-5.0** |  | 9 | **G179**, A208, N209, Q210, T237, D239, F251, V252, **P275**, P277, M278, Y281, A282 |  |
| **Experiment 3** | | | | |
| **Binding energy (kcal/mol)** | **Cluster** | **Pose** | **C*b*_5_R’s residues forming the binding site** | **H-bonds**  **C*b*_5_R-CoQ_0_** |
| **-5.8** | **1** | **1** | **FAD, K110, Y112, G179, G180, T181, G182,** I183**, C273, G274, P275, P276** | **G180-O1** |
| **-5.7** |  | **2** | **FAD, K110, Y112, G179, G180, T181, G182, C273, G274, P275,P276** | **T181-O2** |
| **-5.3** |  | **3** | **FAD, K110, Y112, G179, G180, T181, G182, C273, G274, P275,P276** | **FAD O3B-O1** |
| **-5.2** |  | **4** | **FAD, K110, Y112, G179, G180, T181, G182, C273, G274, P275,P276** |  |
| **-5.1** |  | **5** | **FAD, K110, Y112, G179, G180, T181, G182, C273, G274, P275,P276** |  |
| **-5.0** |  | **6** | **FAD, K110, Y112, G179, G180, T181, G182, C273, G274, P275,P276** |  |
| **-5.0** |  | **8** | **FAD, K110, Y112, G179, G180, T181, G182, C273, G274, P275,P276** | **Y112-O1** |
| -5.0 |  | 7 | **FAD**, H54, D55, **K110**, V111, **Y112**, F113, T116, Q210, D214 |  |
| -4.8 |  | 9 | **FAD,** H54, D55, **K110,** V111**, Y112,** F113, T116, Q210 | Y112-O1 |

Supp. Table 1. Molecular docking results of experiments performed by triplicate are shown in this table. Top 9 ranked poses with individual calculated binding energy, binding site (amino acid residues of Cb5R’s locating within 5 Angstroms from CoQ_0_ moiety) and the H-bonds are shown. Poses were grouped in same clusters in order to define the binding site.
